# Detecting Clinically Relevant Topological Structures in Multiplexed Spatial Proteomics Imaging Using TopKAT

**DOI:** 10.1101/2024.12.18.628976

**Authors:** Sarah Samorodnitsky, Katie Campbell, Amarise Little, Wodan Ling, Ni Zhao, Yen-Chi Chen, Michael C. Wu

## Abstract

Novel multiplexed spatial proteomics imaging platforms expose the spatial architecture of cells in the tumor microenvironment (TME). The diverse cell population in the TME, including its spatial context, has been shown to have important clinical implications, correlating with disease prognosis and treatment response. The accelerating implementation of spatial proteomic technologies motivates new statistical models to test if cell-level images associate with patient-level endpoints. Few existing methods can robustly characterize the geometry of the spatial arrangement of cells and also yield both a valid and powerful test for association with patient-level outcomes. We propose a topology-based approach that combines persistent homology with kernel testing to determine if topological structures created by cells predict continuous, binary, or survival clinical endpoints. We term our method TopKAT (Topological Kernel Association Test) and show that it can be more powerful than statistical tests grounded in the spatial point process model, particularly when cells arise along the boundary of a ring. We demonstrate the properties of TopKAT through simulation studies and apply it to two studies of triple negative breast cancer where we show that TopKAT recovers clinically relevant topological structures in the spatial distribution of immune and tumor cells.

## 1 Introduction

Use of spatially-resolved, multiplexed proteomics imaging is rapidly expanding in oncology research [1, 2, 3, 4]. The ability of spatial proteomic technologies to comprehensively probe the intricate spatial arrangement of tumor, immune, and stromal cells in the tumor microenvironment (TME) provides critical knowledge on how the TME affects tumorigenesis and disease progression [5, 6], treatment response [7], and eventual patient survival [8, 9]. Multiplexed spatial proteomics has been successfully implemented to elucidate the effect of the TME on clinical outcomes in breast [8, 7, 10], colorectal [11], and lung cancers [12, 13], among other diseases [14, 15].

Despite the growing usage of spatial proteomic technologies [16], tools for analyzing the resulting cell-level data lag far behind technical developments, with a serious dearth of tailored, flexible, and robust analytic methods for studying how cellular arrangements are related to patient-level outcomes. Many current findings have been derived from inconsistent, informal or qualitative, or even invalid statistical approaches. Studies using formal, inferential statistical analyses have focused on summarizing the spatial arrangement of cells within a circle of radius *r* using spatial summary statistics [17, 18, 19, 20]. These summary statistics, such as Ripley’s K [21], quantify the degree to which the cells exhibit clustering, dispersion, or complete spatial randomness. This neglects the complex geometric arrangements cells form, for example, around dense tumor regions, necrotic areas, or blood vessels. This geometric information may be clinically or biologically relevant and may be difficult to detect using summary statistics like Ripley’s K. In addition, these methods rely on the assumption of homogeneity or the idea that the number of cells per unit area is constant irrespective of location. If this assumption is violated, as it often is due to gaps or tears in the tissue during sample processing [22], we may lose power to detect an effect on clinical outcomes. Finally, existing approaches are also restricted to a single clinical-outcome type. Several recent approaches work for binary phenotypes (e.g. treatment response) [23, 20, 24] but do not accommodate other important outcomes such as right-censored survival or quantitative outcomes. The lack of appropriate and adaptable tools poses a serious challenge, limiting full utilization and interpretation of the rich data arising in spatial proteomic studies.

The goal of our work is to characterize the geometry of how cells organize in tissue and examine if this geometric characterization associates with patient-level outcomes, such as survival and treatment response. To identify clinically relevant geometric structures among cells, we combine the topological data analysis (TDA) technique known as persistent homology (PH) [25] with kernel association testing to produce the Topological Kernel Association Test (TopKAT). Operationally, TopKAT uses PH to characterize the size and number of homologies, namely connected components and loops, formed by cells in tissue. Then, TopKAT compares whether samples with similar topological or geometric cell structures also exhibit similar clinical phenotypes using kernel association testing. While TDA has seen some basic utility in clinical omics [26, 27, 28, 29], it is notably absent from mainstream analyses of spatial proteomics and existing deployments have not been integrated with formal statistical hypothesis testing. To address this, we integrate PH with the powerful nonparametric kernel association testing framework commonly used for other genomic data types [30, 31, 32].

TopKAT contributes to both statistical methodology and to our ability to derive insights from spatially-resolved, cell-level imaging data, particularly multiplexed spatial proteomics. In terms of statistical methodology, TopKAT combines PH, which is often used descriptively [33], with kernel testing to allow inferential analysis based on the PH summary statistic, the persistence diagram [25], within a unified framework. TopKAT also facilitates deeper clinical and biological insights from spatially-resolved, cell-level imaging through a number of advantages. First, in contrast to existing approaches, leveraging TDA focuses on how cells form geometric structures, including loops, rather than the relationships between pairs of cells. Second, PH detects these structures across a range of scales, offering a holistic assessment of how cells are organized. These advantages allow us to compare different samples on the basis of the presence, absence, and size of geometric features, revealing which samples are, geometrically-speaking, more similar. Third, PH possesses a “smoothing” effect, in the sense that it mitigates noise, such as mislabeled cells or technical artifacts, within images of the heterogeneous biopsies. PH thus produces a rich, yet robust, description of how cells are arranged in tissue. Finally, integration with kernel approaches allows one to utilize these frameworks for rigorous and robust statistical inference across various study designs, clinical outcome types, and analytic objectives. We demonstrate through extensive simulation studies that TopKAT offers higher power and better type I error control than existing alternatives which utilize spatial summary statistics. Further, we apply TopKAT to data from two studies of triple negative breast cancer (TNBC) [8, 7] to illustrate the importance of topological structure among cells in describing patient survival and response to immunotherapy treatment.

## 2 Results

### 2.1 Overview of TopKAT

TopKAT is a global test for the association between topological (geometric) structures within the spatial distribution of cells in cell-level imaging and sample-level outcomes, adjusting for potential confounders. The motivation is that the geometry of the spatial distribution of cells may characterize a phenotype with clinical implications.

Operationally, TopKAT follows a three-step procedure (Figure 1). First, TopKAT captures the topological structure of the cells within each image using PH and by applying a Rips filtration [33]. PH reveals the abundance and size of homologies, the generalized concept of a hole. Since our single-cell images are two-dimensional, we are interested in detecting connected components (which represent dense masses of cells) and loops (which represent regions where cells arise along the boundary of a ring but are absent from the center.) The topological structure of each image is characterized in a summary statistic termed a persistence diagram [33].

**Figure 1:**
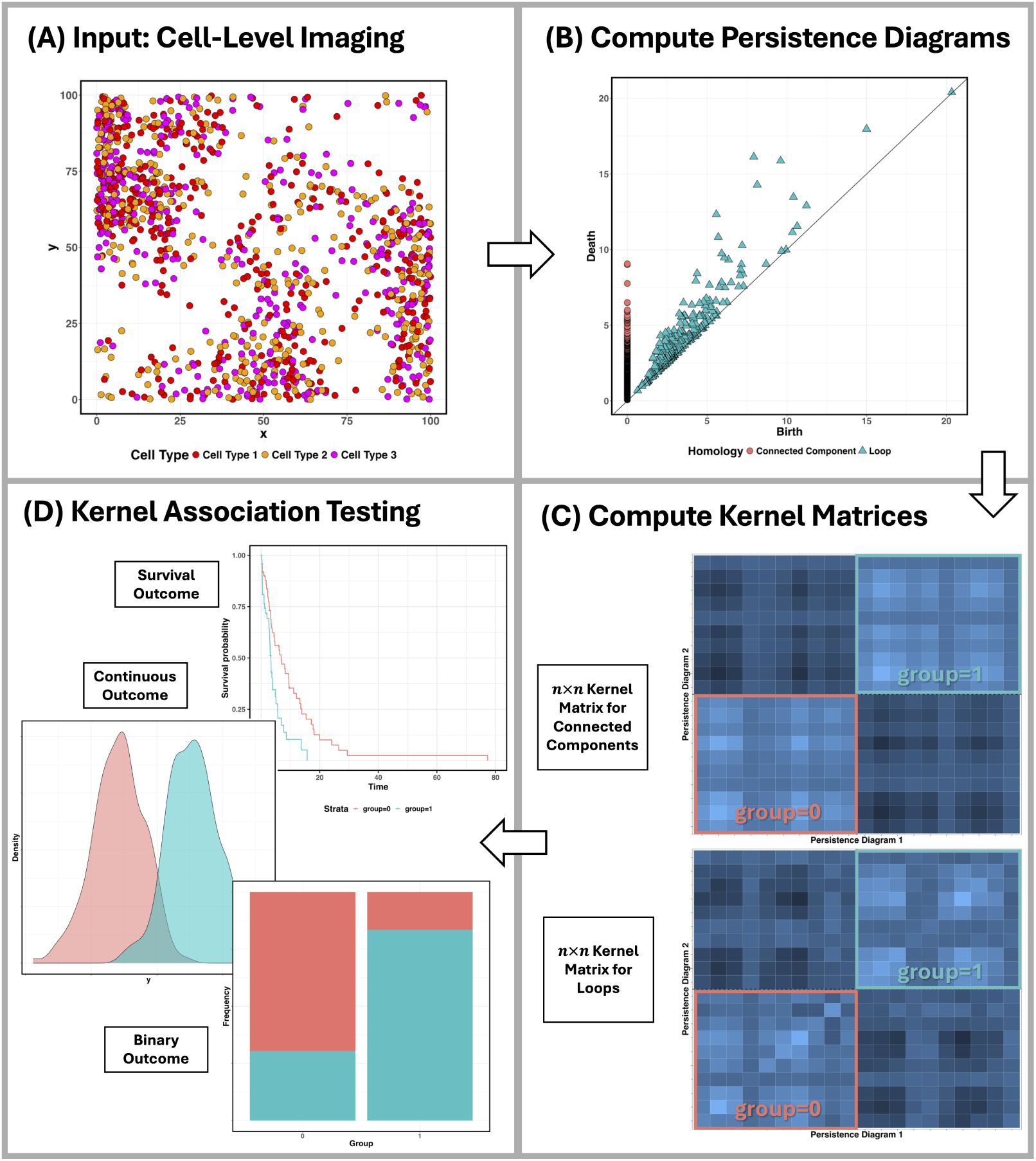
Flow chart illustrates the steps of the TopKAT method. The input data for TopKAT are cell-level images (A). TopKAT then characterizes the topological structures created by cells in each image and summarizes this information via persistence diagrams (B). TopKAT then calculates the similarity between each pair of persistence diagrams, quantified in kernel matrices where each entry reflects how similarity between corresponding pairs of persistence diagrams (C). In this example, the images arise from two groups of patients (group 0 and group 1) whose cell-level images exhibit different topological structures. The similarities between images within these groups are highlighted in red and blue boxes. TopKAT then inputs the kernel matrices into a kernel machine regression model to test for an association with clinical outcomes (D).

The second step is to compute a pairwise distance matrix quantifying the distance (or dissimilarity) between each pair of persistence diagrams. We construct a distance matrix for each homology (connected components and loops) which allows us to describe the differences between samples based on each type of topological structure. We then convert these distance matrices to kernel matrices which describe the pairwise similarity between persistence diagrams.

Finally, we use a kernel testing framework [34] to test for an association between the similarity in persistence diagrams and similarities in continuous, binary, or survival outcomes, adjusting for covariates. To include information from each homology group, we consider a series of weighted combinations of the kernel matrices for the connected components and loops. We iterate across each potential combination and aggregate the resulting p-values using the Cauchy combination test [35]. If similarities among the persistence diagrams (quantity and size of homologies) align with similarities in the outcome, topological structures within the TME are associated with outcomes, such as treatment response or survival.

There are several advantages to using TopKAT. First, PH describes the geometry of the spatial distribution of cells which can be used to compare the topological differences between samples with distinct outcomes. This description of the spatial distribution of cells is robust to perturbations in the data due to technical artifacts [33]. This is essential because our analysis is downstream of a complex normalization, segmentation, and phenotyping pipeline which may contribute to noise in the images, such as mislabeled cells. Second, PH can be used to identify global structures within the TME, such as large masses of immune cells or regions within the TME where immune cells are unable to penetrate. These attributes are difficult to capture by other approaches, such as spatial summary statistics [22]. These global structures have been shown to be clinically informative in breast cancer [8, 7], which we explore in Sections 2.3 and in the Supplementary Materials Section 7.

### 2.2 Simulation Study

We evaluated the validity (type I error control) and power of TopKAT on simulated images and compared it to existing approaches to assess the advantage of using topological information to predict clinical outcomes. We studied three variations of TopKAT: in the first, we only considered similarities in the number and size of connected components across images; in the second, we only considered similarities among loops; and in the third, we aggregated the kernel matrices across similarities in both homology groups using the Cauchy combination test (Supplementary Materials Section 2). We compared TopKAT to existing approaches grounded in the spatial point process model, including SPOT [17], FunSpace [18], and SPF [19], which we henceforth refer to as “spatial methods.” These methods describe the spatial arrangement of cells within circles of radius *r*, varying the size of this radius.

We simulated *n* = 100 samples exhibiting different geometric structures among the cells. We split the samples into two groups of 50 and selected a different geometric structure to randomly simulate within each group. This included random numbers of squares (“square”), loops (“loop”), bivariate Gaussian distributions (“clusters”), simulated tissue images using the scSpatialSIM R package [36] (“simulated tissue”), and complete spatial randomness (“CSR”). Examples of these images are provided in the Supplementary Materials Section 5.2. The simulation conditions are referred to based on the geometric structures chosen for each group, e.g., squares vs. loops, loop vs. CSR. To assess power, survival outcomes for these two groups were simulated from exponential distributions with different rates with a hazard ratio of 2. To assess validity, the outcomes across both groups were simulated from the same exponential distribution. We simulated survival outcomes, though TopKAT accommodates continuous and binary outcomes which are illustrated in our data applications.

The results are given in Table 1. Across all conditions, TopKAT offers higher power, and across most conditions, more tightly controls type I error than the spatial approaches. This suggests that (1) topological information (connected components and loops) captures the different geometric and spatial arrangements among the cells and (2) our choice of distance measure and kernel function capture the similarities and differences in this topological information between images. TopKAT far exceeds the spatial methods in power under scenarios in which loops were simulated (“loop vs. CSR” or “square vs. loop”). All three variations of TopKAT exhibited power around 0.89, whereas the spatial methods exhibited power between 0.52 and 0.69. This is because TopKAT explicitly detects the presence and size of loops while the spatial methods capture cell density within a specific radius. TopKAT, however, also detects differences in cell density, as shown by the “square vs. CSR” and “cluster vs. CSR” conditions. Here, the images contained either tight masses of cells (squares, which reflect large connected components), diffuse masses of cells (clusters, which also reflect connected components), or random noise (CSR). TopKAT and the spatial methods showed comparable power (around 0.88 for TopKAT and between 0.78 and 0.85 for the spatial methods). Finally, TopKAT is sensitive to detecting differences due to tissue structure. Under “simulated tissue,” TopKAT detected differences in tissue structure were associated with sample-level phenotypes. Identifying these structural associations is essential for real applications [29].

**Table 1:** Power and type I error rates for TopKAT vs. spatial methods (SPOT, SPF, FunSpace) across a range of simulated images for a survival outcome. TopKAT (dim 0) refers to TopKAT using only the kernel matrix derived from the persistence of dimension-0 homologies (connected components). TopKAT (dim 1) refers to TopKAT using the kernel matrix derived from persistence of dimension-1 homologies (loops).

| Model | square vs. CSR |  | loop vs. CSR |  | square vs. loop |  | cluster vs. CSR |  | simulated tissue |  |
| --- | --- | --- | --- | --- | --- | --- | --- | --- | --- | --- |
|  | Power | Type I Error | Power | Type I Error | Power | Type I Error | Power | Type I Error | Power | Type I Error |
| TopKAT (dim 0) | 0.899 | 0.044 | 0.894 | 0.054 | 0.830 | 0.054 | 0.886 | 0.048 | 0.877 | 0.050 |
| TopKAT (dim 1) | 0.882 | 0.050 | 0.895 | 0.060 | 0.863 | 0.059 | 0.885 | 0.045 | 0.876 | 0.052 |
| TopKAT (omnibus) | 0.907 | 0.048 | 0.895 | 0.058 | 0.851 | 0.055 | 0.884 | 0.047 | 0.878 | 0.052 |
| SPOT | 0.770 | 0.051 | 0.610 | 0.055 | 0.593 | 0.052 | 0.845 | 0.052 | 0.800 | 0.059 |
| SPF | 0.754 | 0.059 | 0.609 | 0.063 | 0.685 | 0.067 | 0.781 | 0.058 | 0.726 | 0.069 |
| FunSpace | 0.681 | 0.047 | 0.516 | 0.054 | 0.578 | 0.058 | 0.825 | 0.050 | 0.669 | 0.059 |

Within TopKAT, we compared using topological information from each homology group: only the connected components (“dim 0” in Table 1) or only the loops (“dim 1” in Table 1). We contrasted these with an omnibus test across both groups (“omnibus” in Table 1). Intuitively, in scenarios with dense regions of cells (e.g. “square vs. CSR”), testing solely based on similarities among the connected components should yield more power. We observed this to be the case, as TopKAT (dim 0) power (0.899) slightly exceeded the power of TopKAT (dim 1) (0.882) in this condition. In scenarios where loops were simulated, detecting only loops should be more powerful than detecting only connected components. We found that this was the case, though the gain in power was slight (0.895 for TopKAT (dim 1) vs. 0.894 for TopKAT (dim 0)). Overall, neglecting the “correct” topological structure for each scenario did not lead to a marked reduction in power. For example, using only the loops yielded power of 0.88 (vs. 0.90 for TopKAT (dim 0)) under “square vs. CSR.” This reflects the subtle topological information available in these images. For example, in images simulated under CSR, there are small loops present among the randomly-scattered cells which distinguishes these images from large square masses. In general, however, TopKAT using an omnibus approach offered similar or higher power compared with using a single homology group.

### 2.3 Application to Multiplexed Ion Beam Imaging of Triple Negative Breast Cancer

We applied TopKAT to multiplexed ion beam imaging time-of-flight (MIBI-TOF) data obtained from a study of triple negative breast cancer (TNBC) [8]. This study used MIBI-TOF to probe the spatial expression of 36 proteins in breast tumor tissue biopsied from 38 TNBC patients. The goal of the study was to explore the associations between the cellular structure of the TME and clinical endpoints, including overall survival. The data contains images selected as representative of the cellular structure within each tumor sample. Cells were phenotyped as either immune cells (CD4 T cells, CD8 T cells, CD3 T cells, natural killer cells, B cells, macrophages, dendritic cells, neutrophils, or monocytes) or non-immune cells (tumor cells, epithelial cells, mesenchymal cells, or endothelial cells).

Keren et al. discovered that the biopsies exhibited distinct “structured” TMEs. Biopsies could be categorized as either (1) compartmentalized, where immune and tumor cells segregated into distinct regions from each other (Figure 2A), (2) mixed, where immune and tumor cells colocalized (Figure 2B), or (3) cold, where few immune cells were detected within the tumor compartment of the biopsy. These categories further exhibited differential survival outcomes, with compartmentalization associated with improved survival, underscoring the clinical relevance of these global cell structures within the TME. Of the *n* = 38 samples, 18 exhibited mixed TMEs, 15 exhibited compartmentalization, and 5 were immune cold. These categories were determined based on a “mixing score” which was computed as the number of immune-tumor cell interactions divided by the number of immune-immune cell interactions. The resulting categories (mixed, compartmentalized, and cold) were provided in the published data. The goals of our analysis were to: (1) characterize the global structure of immune cells among the biopsies using persistent homology and (2) relate this structure to overall survival.

**Figure 2:**
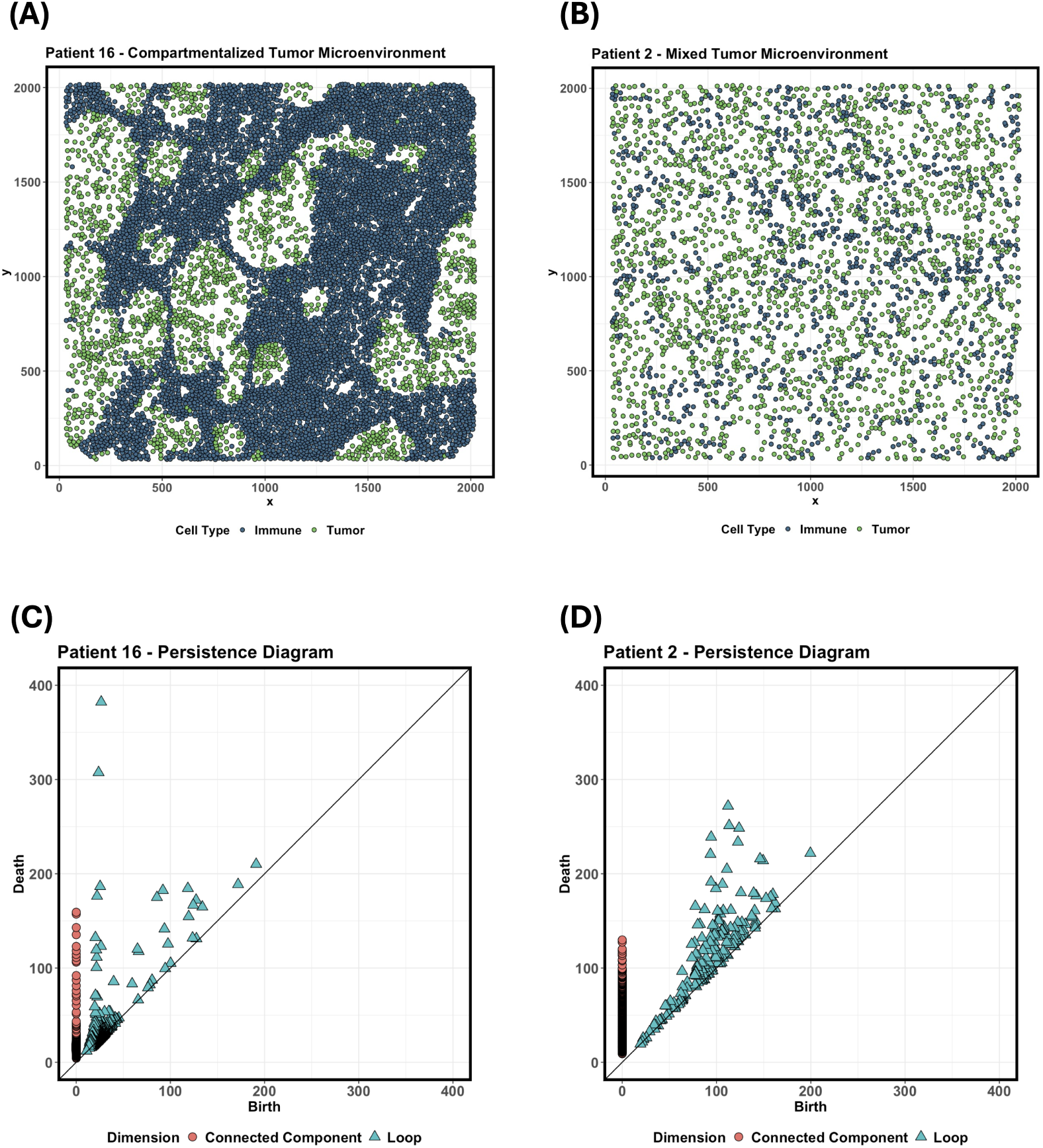
Comparing the persistence diagrams for two example mixed and compartmentalized tumor microenvironments. Examples of compartmentalized (A) and mixed (B) samples show distinct spatial arrangements of tumor and immune cells. Persistence diagrams illustrating the scales and lifespans of connected components and loops among immune cells are shown in (C) and (D) for the example compartmentalized TME and example mixed TME. In the compartmentalized TME, connected components and loops among immune cells tend to persist longer than the mixed TME. Loops also tend to be larger in compartmentalized TMEs, as evidence by their longer lifespans.

In the Supplementary Materials Section 6, we describe how TopKAT significantly differentiates between mixed, compartmentalized, and cold TMEs. Here, we focus on mixed and compartmentalized samples and closely examine the topological structures inherent to immune cells among these biopsies. Figures 2C and 2D show persistence diagrams in two example compartmentalized and mixed samples. Comparing these two, the compartmentalized TME contained connected components and loops that persisted longer than in the mixed TME. Indeed, the average maximum lifespan for connected components in compartmentalized TMEs was slightly larger than in mixed TMEs (179.69 vs. 171.66) and was much larger for loops (215.29 vs. 164.45). This is intuitive: compartmentalized TMEs exhibit large masses of immune cells which persist longer as connected components within the Rips filtration than in mixed TMEs. Compartmentalized TMEs may also show large gaps or loops representing regions where tumor cells resided that prevented immune cells from infiltrating.

We then followed the procedure given in Section 7.5 to identify the distance between cells at which mixed and compartmentalized TMEs showed the most distinctions in topological structure. TopKAT yielded the smallest p-value at a distance of 73.6 (Supplementary Figure 2). In Figures 3A and 3B we illustrate the simplicial complex built based on the immune cells at this distance on two example samples. These figures show a close recapitulation of the immune cell structures visualized in Figures 2A and 2B. We then explored how often immune cells were connected at a distance of 73.6. On average, compartmentalized TMEs exhibited a large number of connections within B cells, CD4 T cells, macrophages, and CD8 T cells (Figure 3C). These connections potentially reflect the presence of tertiary lymphoid structures, which arise at sites of consistent inflammation [37]. On average, mixed TMEs exhibited connections among these immune cell types, but also included more diverse connections among CD3 T cells, T regulatory cells (Treg), and neutrophils (Figure 3D).

**Figure 3:**
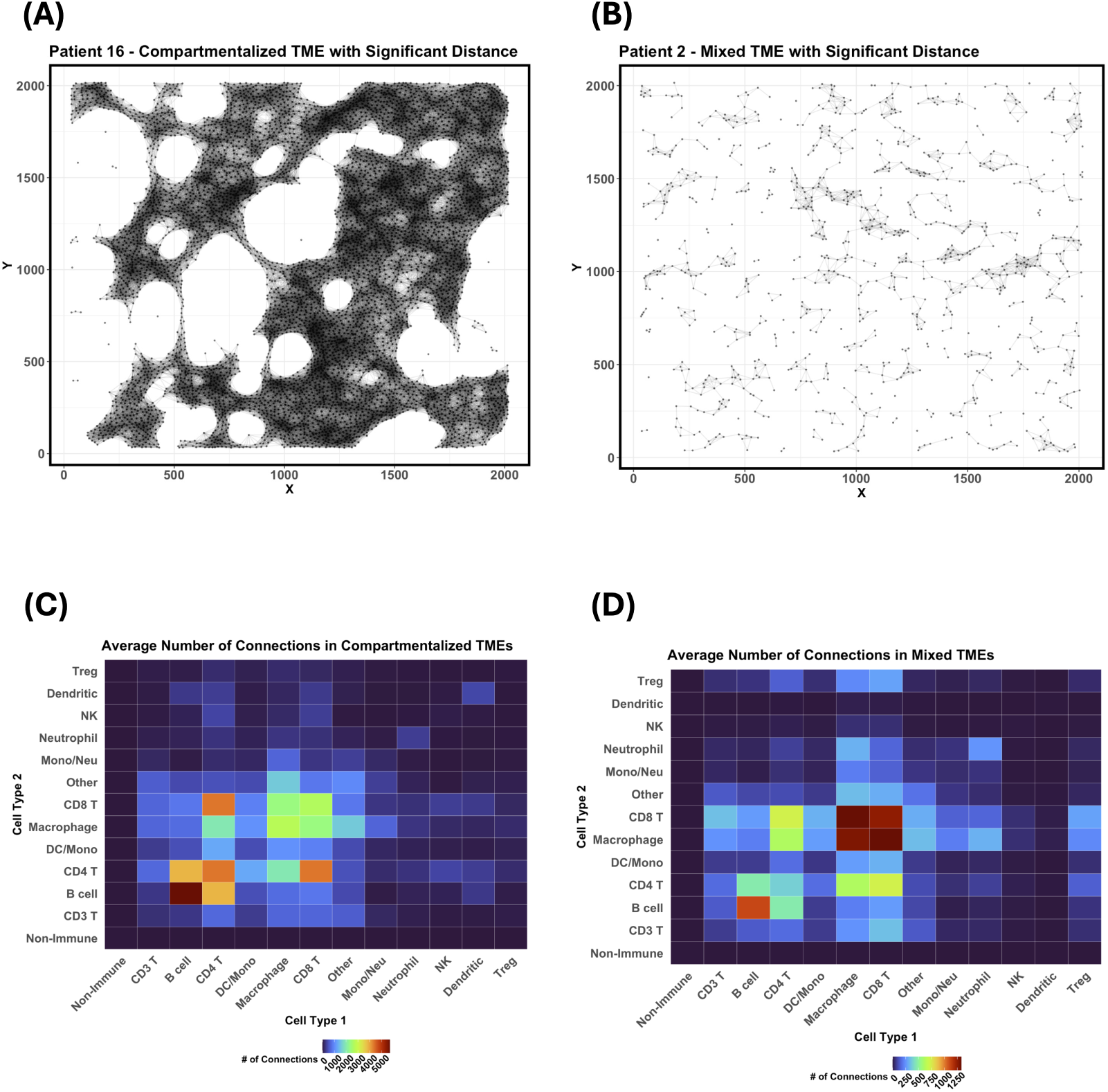
Exploring the connections among immune cells at a clinically relevant distance. We obtained the distance at which mixed and compartmentalized tumor microenvironments (TMEs) are most distinct topologically, which occurs at distance 2*ɛ* = 73.6. Two example compartmentalized (A) and mixed (B) samples with this distance overlaid are shown. We then illustrate the average number of connections between immune cell phenotypes at this distance in Figures (C) and (D) among compartmentalized and mixed TMEs, respectively.

Finally, we examined if similarities in the topology of each TME predicted overall survival. Consistent with the original study findings, we observed mixed, segregated, and cold TMEs corresponded to distinct survival outcomes (*p* = 0.011). Note that the spatial methods considered in Section 2.2 (SPOT, SPF, and FunSpace) could not consistently recapitulate these results. For these approaches, we considered a range of radii between 0 and 512. We did not find significant associations between the spatial arrangement of immune cells and survival using these approaches. Only SPOT and FunSpace were able to detect pairwise differences in immune cell structure between mixed and cold TMEs (both methods) and compartmentalized and cold TMEs (FunSpace only).

In the Supplementary Materials Section 7, we describe using TopKAT to analyze an additional study of triple negative breast cancer to examine how the TME differs between responders and non-responders to immunotherapy treatment.

## 3 Discussion

Imaging with cell-level resolution, including multiplexed spatial proteomics, can reveal the cellular composition of the tumor microenvironment which is known to associate with clinical outcomes. There is an absence of valid and robust statistical methods for testing for this association and a lack of consistency among methods across studies. To address this gap, we combined persistent homology and kernel association testing in TopKAT. TopKAT is a global test of whether similarities in the quantity and scale of homologies among the single-cell images aligns with similarities in a clinical outcome to determine if topological structure among the cells is clinically informative.

We illustrated the power and type I error rate of TopKAT on simulated data and found that TopKAT outperforms existing methods for revealing the relationship between cell structures in the image and clinical outcomes. This was especially apparent when the cell architecture contained loops which persistent homology is well-suited to capture. TopKAT may be modified to up- or down-weight the contribution of each degree-*k* homology to suit the hypotheses of the investigator. In our simulations, we found that aggregating across connected components and loops offered similar or higher power than considering one dimension alone. This may be particularly beneficial in scenarios where the importance of each homology is unknown. However, if *a priori* knowledge suggests a certain homology may be more strongly associated with sample-level outcomes, the weighted aggregation of the kernel matrices may be modified to suit. An example scenario where this could arise is in samples known to exhibit regions of necrosis, where immune cells may not be able to infiltrate.

There are several limitations and directions to expand upon this work. First, TopKAT does not distinguish between cell types in construction of the filtration. In our two triple negative breast cancer applications, we used TopKAT to examine the spatial architecture of cells irrespective of phenotype. After the fact, we examined which immune cell types were highly connected within the simplicial complex formed at a clinically-relevant distance between cells. Examining these connections may reveal important structures among distinct cell phenotypes. However, the interpretation based on the TopKAT p-value is still agnostic to cell type. TopKAT could be run on each cell type separately, or on any subset of the cell types, though this would not distinguish the contributions of each cell type in constructing the homologies. We also presented TopKAT based on the construction of a Rips filtration, the use of a total dissimilarity based the lifespans of homologies, and a Gower’s centered kernel. These choices yielded a fast and intuitive test, but could be modified. Different distances or dissimilarities may uncover more pronounced differences among the persistence diagrams. Other kernels, such as an exponential kernel, may better capture the relationship between topological structure of the cells and patient outcomes. The impact of these choices is beyond the scope of this article, but warrants further evaluation. Another limitation is that this method does not accommodate multiple regions-of-interest imaged within the same biopsy. This could be addressed by considering an aggregation of the persistence diagrams across images within a biopsy, but this remains to be explored in future work. Finally, TopKAT is a global test of association between the topological structure of the images and outcomes. One could consider, say, examining if the average lifespan of a particular homology, such as the connected components, associates with patient outcomes and estimate the effect of an increase in the average lifespan on the outcome.

## 4 Data Availability

The data for the application given in Section 2.3 can be found at https://www.angelolab.com/ mibi-data. The data for the NeoTRIP application given in the Supplementary Materials Section 7 can be found at https://zenodo.org/records/7990870.

## 5 Code Availability

Software to implement TopKAT is available in an R package available at https://sarahsamorodnitsky. github.io/TopKAT/.

## Acknowledgements

This work was supported in part by NIH Grant U10 CA180819 and The Hope Foundation for Cancer Research.

## 6 Methods

### 6.1 Notation

Assume we have *n* two-dimensional multiplexed spatial proteomics images obtained from tissue biopsies collected from *n* individuals. In our case, we are interested in examining the spatial distribution of cells in the tumor microenvironment and use language pertaining this application. However, this methodology is appropriate for any study involving the spatial arrangement of cells in tissue. Each image is represented by a matrix of 2D (*x, y*) coordinates for the location of each cell. Let **y** : *n* × 1 represent a vector of patient-level outcomes, which may be continuous, binary, or right-censored survival endpoints. Let **X** : *n* × *p* represent a matrix of *p* clinical covariates, e.g. sex or age, that we would like to adjust for in our kernel association test.

Our goal is to test for an association between the topological features within each image and clinical outcomes, adjusting for covariates. To achieve this, we proceed in the following steps:

1. We first summarize the topological information contained in each image using persistence diagrams. We obtain these diagrams using persistent homology and by constructing a Vietoris-Rips filtration [25].
2. We then construct an *n* × *n* pairwise distance matrix quantifying the distance between each pair of persistence diagrams. We obtain a distance matrix for each homology group (connected components and loops). To facilitate the use of kernel machine regression, we convert the distance matrices to kernel matrices which quantify the similarity between persistence diagrams for each homology.
3. Finally, we use kernel machine regression to relate the similarity in persistence diagrams to clinical outcomes, adjusted for covariates. We describe these steps in more detail below.

### 6.2 Computing Persistent Homology

The first step in the TopKAT pipeline is to summarize the topological information in each image using persistent homology. Persistent homology is a topological data analysis technique that captures the size of connected components (homologies of degree-0) and loops (homologies of degree-1) within a 2D point cloud [33]. To capture these features, persistent homology relies on constructing an evolving graph of nodes and vertices where the nodes represent each point in the point cloud (in our case, cells in an image). The graph, known as a *simplicial complex*, evolves based on a scale parameter *ɛ* that is varied from 0 to ∞. As *ɛ* changes, connected components (homologies of degree-0) and loops (homologies of degree-1) will appear (birth) and disappear (death) in the resulting graph and the difference between the birth and death scales is termed the *lifespan* of a feature. Geometric features with longer lifespans correspond to larger (i.e., “more persistent”) features of the data. The most persistent connected components and loops are the ones we hope to capture and will introduce the most information in our subsequent kernel association test.

The process of varying *ɛ* to generate a sequence of simplicial complexes is called constructing a *filtration*. We compute a Vietoris-Rips filtration (also known as a Rips filtration) because it is fast, efficient, and intuitive. Let *S* represent a 2D spatial image of cells. For *ɛ* ≥ 0, the simplicial complex generated within the Rips filtration is defined as

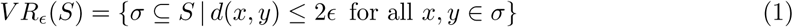

In plain terms, the Rips complex at *ɛ* is a subset, *σ*, of *S* that contains the *ɛ*-neighborhood graph of *S* [33]. As *ɛ* grows, we obtain a sequence of nested subsets, *σ_ɛ_*_1_ *, σ_ɛ_*_2_ *, …* which allows us to study the evolution of connected components and loops in this space.

The values of *ɛ* at which each geometric feature is born and dies is recorded and summarized using a *persistence diagram* [33]. Formally, a persistence diagram, *Z*, is a multiset of points defined as

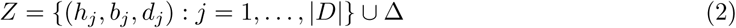

where *h_j_* represents the homology degree of the *j*th point, *b_j_* represents the scale at which the *j*th feature is born, and *d_j_* represents the scale at which the *j*th feature dies. |*D*| counts the total number of connected components and loops detected during filtration and Δ refers to the all the points along the diagonal *y* = *x* line [38]. We will use the persistence diagram as our summary statistic to capture the geometric information about the spatial distribution of cells in each multiplexed image. Computation of persistent homology is efficiently implemented in the TDAstats R package [39].

### 6.3 Distance and Kernel Matrix Construction

Given a sequence of persistence diagrams, **Z** = (*Z*_1_*, …, Z_n_*), we would like to use these as covariates in a test of association between topological structure in our images and clinical outcomes. To do this, we use a kernel machine learning framework which relies on first constructing a kernel matrix, **K**. To obtain **K**, which describes the similarity between persistence diagrams, we first construct a pairwise distance matrix between the samples and convert this to a kernel matrix. Since persistent homology emphasizes uncovering “persistent” degree-0 and degree-1 homologies, we use the following dissimilarity measure which we term *total dissimilarity* :

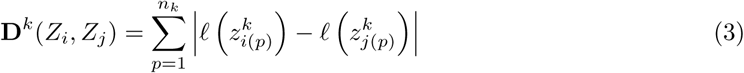

where *n^k^*= max(*n_i_^k^, n_j_^k^*) is the maximum number of homologies of degree-*k* between *Z_i_* and *Z_k_* and 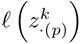 is the lifespan of the *p*th longest-living homology of degree *k* in persistence diagram *Z_j_*.

To compute this distance, the features of degree-*k* are first ordered from highest to lowest lifespan. Any difference in the number of features is filled in with zeros. The sum of deviations between ordered lifespan statistics is computed as given in Equation 3. **D***^k^*(*Z_i_, Z_j_*) emphasizes a similarity in (1) quantity and (2) lifespan of degree-*k* homologies found in persistence diagrams *Z_i_* and *Z_j_*.

Intuitively, *Z_i_* and *Z_j_* with a similar number of degree-*k* features with similar lifespans will have a comparatively small **D***^k^*(*Z_i_, Z_j_*). This yields *K* distance matrices corresponding to each of the degree-*k* homologies, *k* = 1*, …, K*. These distance matrices, denoted **D***^k^*, are converted to kernel matrices using a Gower’s centered kernel:

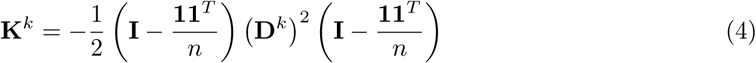

where **I** : *n* × *n* is the identity matrix and **1** is a *n* × 1 column vector of 1s. We convert the **D***^k^* to kernel matrices to leverage kernel machine regression, which we discuss in the next section.

### 6.4 Kernel Machine Regression

We now would like to relate the persistence diagrams, **Z**, to a clinical outcome, **y**, via a kernel machine regression model. Here we focus on a survival outcome, but this framework extends to continuous and binary outcomes which are discussed in the Supplementary Materials. For a survival outcome, suppose we observe 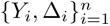 where *Y_i_* = min(*T_i_, C_i_*) is the recorded event time (either the time of the survival event *T_i_* or the censoring time *C_i_*) and Δ*_i_* = *I*(*T_i_* ≤ *C_i_*) is an event indicator. To test if **Z** is associated with survival time, we use a kernel machine Cox proportional hazards model:

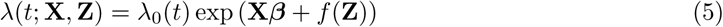

where ***β*** represents the effect of known confounders on the log-hazard for survival and *f* (·) is a smooth, unknown, and centered vector-valued function that is assumed to belong to a space spanned by a positive-definite kernel *K*(·, ·) [34]. The kernel function, *K*(·, ·), quantifies the similarities between samples *Z_i_* and *Z_j_, i*, ≠ *j* which, in our case, are persistence diagrams. We are interested in testing if *f* (**Z**) has an effect on the log-hazard for survival. Intuitively, if similarities between persistence diagrams aligns with similarities in outcomes, we should expect to see a non-zero effect of *f* (**Z**) on **y** [31].

Within this kernel machine Cox regression model, the null and alternative hypotheses we wish to test are:

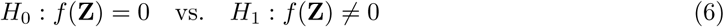

To test *H*_0_, we leverage the relationship between kernel machine Cox regression and linear mixed modeling [40]. We can rewrite the model given in Equation 5 as:

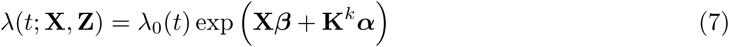

where (***β****, **α***) are unknown parameters to be estimated. Our null hypothesis then becomes *H*_0_ : **K***^k^**α*** = 0. Maximizing the penalized partial likelihood corresponding to Equation 7 is equivalent to estimating a Cox survival model with a random intercept for each sample:

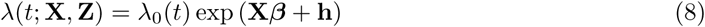

where **h***^T^* = (*h*_1_*, …, h_n_*)*^T^* ∼ Normal(**0***, τ* **K***^k^*) is a vector of random effects with covariance proportional to the kernel matrix for homology degree-*k*, **K***^k^*. Therefore, testing *H*_0_ : **K***^k^**α*** = 0 is the same as testing *H*_0_ : *τ* = 0. We can test this null hypothesis using the variance-component score test which requires only fitting the null model under *τ* = 0, *λ*(*t*; **X**, **Z**) = *λ*_0_(*t*) exp (**X*β***). This implies that the test is a valid statistical test even if an undesirable kernel is chosen. However, the choice of kernel will impact power [31, 41]. An exploration of the properties of a desirable kernel are beyond the scope of this work.

This variance-component score test statistic for the kernel machine Cox model based on degree-*k* is:

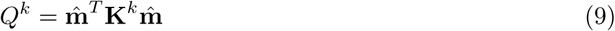

where **m̂** = (*m̂* _1_*, …, m̂* *_n_*) represents a vector of Martingale residuals under *H*_0_ where 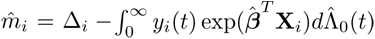 and 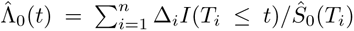 is Breslow’s estimator of the baseline hazard function under the null and *Ŝ*_0_(*t*) is the estimator of the baseline survival function. To handle tied survival times, we use Efron’s approximation [42]. More information is given in [41]

Under *H*_0_, *Q^k^* has an asymptotic distribution of a mixture of *χ*_1_^2^ distributions. The p-value can then be computed analytically using Davies method [43].

In the Supplementary Materials Sections 1 and 2, we discuss a small sample correction and accommodating kernel matrices from both connected components and loops. Both approaches were used in the TNBC application described in Section 2.3. We also discuss adaptations of TopKAT for continuous and binary outcomes in Supplementary Materials Sections 3 and 4.

### 6.5 Interpretation

Persistent homology examines the longevity of topological features in each image throughout the filtration. We consider a stepwise approach to determining the distance between cells, 2*ɛ*, within the Rips filtration where we observe the strongest association between persistence diagrams and outcomes.

We first select the maximum distance at which a death of a topological feature was observed across all *n* persistence diagrams. We call this distance 2*ɛ_max_*. We then consider a sequence of length *T* of distances between 0 and 2*ɛ_max_*. For each *ɛ_t_* for *t* = 0*, …, T* (*ɛ*_0_ = 0), we threshold the persistence diagrams to only features which were born and had died during filtration by distance 2*ɛ_t_*. We perform the kernel machine regression test described in Section 7.4 and store the resulting omnibus p-value. At the end, we identify which distance yielded the smallest p-value. Denote the selected distance 2*ɛ_best_*. 2*ɛ_best_* is then interpreted as the maximum distance between cells at which the lifespans and number of topological features was most strongly associated with the outcome. Note that in this case, we are treating the p-value as a statistic and would not recommend interpreting these as p-values in the traditional sense.

Once the “best” distance is obtained, we can conduct further post-hoc analyses. For example, we can consider which cell types are highly connected within the resulting simplicial complex constructed at the identified radius.

## References

[1] Bernd Bodenmiller. Highly multiplexed imaging in the omics era: understanding tissue structures in health and disease. Nature Methods, 21(12):2209–2211, 2024.

[2] Sabrina M Lewis, Marie-Liesse Asselin-Labat, Quan Nguyen, Jean Berthelet, Xiao Tan, Verena C Wimmer, Delphine Merino, Kelly L Rogers, and Shalin H Naik. Spatial omics and multiplexed imaging to explore cancer biology. Nature methods, 18(9):997–1012, 2021.

[3] Yamei Chen, Yuan Liu, and Leng Han. Spatial landscape of the tumor immune microenvironment. Trends in Cancer, 2023.

[4] Emma Lundberg and Georg HH Borner. Spatial proteomics: a powerful discovery tool for cell biology. Nature Reviews Molecular Cell Biology, 20(5):285–302, 2019.

[5] Daniela F Quail and Johanna A Joyce. Microenvironmental regulation of tumor progression and metastasis. Nature medicine, 19(11):1423–1437, 2013.

[6] Karin E de Visser and Johanna A Joyce. The evolving tumor microenvironment: From cancer initiation to metastatic outgrowth. Cancer Cell, 41(3):374–403, 2023.

[7] Xiao Qian Wang, Esther Danenberg, Chiun-Sheng Huang, Daniel Egle, Maurizio Callari, Begoña Bermejo, Matteo Dugo, Claudio Zamagni, Marc Thill, Anton Anton, et al. Spatial predictors of immunotherapy response in triple-negative breast cancer. Nature, 621(7980):868–876, 2023.

[8] Leeat Keren, Marc Bosse, Diana Marquez, Roshan Angoshtari, Samir Jain, Sushama Varma, Soo-Ryum Yang, Allison Kurian, David Van Valen, Robert West, et al. A structured tumor-immune microenvironment in triple negative breast cancer revealed by multiplexed ion beam imaging. Cell, 174(6):1373–1387, 2018.

[9] H Raza Ali, Leon Chlon, Paul DP Pharoah, Florian Markowetz, and Carlos Caldas. Patterns of immune infiltration in breast cancer and their clinical implications: a gene-expression-based retrospective study. PLoS medicine, 13(12):e1002194, 2016.

[10] Esther Danenberg, Helen Bardwell, Vito RT Zanotelli, Elena Provenzano, Suet-Feung Chin, Oscar M Rueda, Andrew Green, Emad Rakha, Samuel Aparicio, Ian O Ellis, et al. Breast tumor microenvironment structures are associated with genomic features and clinical outcome. Nature genetics, 54(5):660–669, 2022.

[11] Christian M Schürch, Salil S Bhate, Graham L Barlow, Darci J Phillips, Luca Noti, Inti Zlobec, Pauline Chu, Sarah Black, Janos Demeter, David R McIlwain, et al. Coordinated cellular neighborhoods orchestrate antitumoral immunity at the colorectal cancer invasive front. Cell, 182(5):1341–1359, 2020.

[12] Hartland W Jackson, Jana R Fischer, Vito RT Zanotelli, H Raza Ali, Robert Mechera, Savas D Soysal, Holger Moch, Simone Muenst, Zsuzsanna Varga, Walter P Weber, et al. The single-cell pathology landscape of breast cancer. Nature, 578(7796):615–620, 2020.

[13] Elham Karimi, Miranda W Yu, Sarah M Maritan, Lucas JM Perus, Morteza Rezanejad, Mark Sorin, Matthew Dankner, Parvaneh Fallah, Samuel Doré, Dongmei Zuo, et al. Single-cell spatial immune landscapes of primary and metastatic brain tumours. Nature, 614(7948):555–563, 2023.

[14] Nicolas Damond, Stefanie Engler, Vito RT Zanotelli, Denis Schapiro, Clive H Wasserfall, Irina Kusmartseva, Harry S Nick, Fabrizio Thorel, Pedro L Herrera, Mark A Atkinson, et al. A map of human type 1 diabetes progression by imaging mass cytometry. Cell metabolism, 29(3):755– 768, 2019.

[15] Sudipa Maity, Yuanyu Huang, Mitchell D Kilgore, Abbigail N Thurmon, Lee O Vaasjo, Maria J Galazo, Xiaojiang Xu, Jing Cao, Xiaoying Wang, Bo Ning, et al. Mapping dynamic molecular changes in hippocampal subregions after traumatic brain injury through spatial proteomics. Clinical Proteomics, 21(1):32, 2024.

16. Method of the Year 2024: spatial proteomics. Nature Methods, 21(12):2195–2196, December 2024. Publisher: Nature Publishing Group.

[17] Sarah N Samorodnitsky, Katie M Campbell, Antoni Ribas, and Michael C Wu. A spatial omnibus test (spot) for spatial proteomic data. bioRxiv, pages 2024–03, 2024.

[18] Thao Vu, Souvik Seal, Tusharkanti Ghosh, Mansooreh Ahmadian, Julia Wrobel, and Debashis Ghosh. Funspace: A functional and spatial analytic approach to cell imaging data using entropy measures. PLOS Computational Biology, 19(9):e1011490, 2023.

[19] Thao Vu, Julia Wrobel, Benjamin G Bitler, Erin L Schenk, Kimberly R Jordan, and Debashis Ghosh. Spf: a spatial and functional data analytic approach to cell imaging data. PLoS computational biology, 18(6):e1009486, 2022.

[20] Nicolas P Canete, Sourish S Iyengar, John T Ormerod, Heeva Baharlou, Andrew N Harman, and Ellis Patrick. spicyr: spatial analysis of in situ cytometry data in r. Bioinformatics, 38(11):3099–3105, 2022.

[21] Brian D Ripley. Modelling spatial patterns. Journal of the Royal Statistical Society: Series B (Methodological), 39(2):172–192, 1977.

[22] Julia Wrobel, Coleman Harris, and Simon Vandekar. Statistical analysis of multiplex immunofluorescence and immunohistochemistry imaging data. Statistical Genomics, pages 141– 168, 2023.

[23] Souvik Seal, Brian Neelon, Peggi M Angel, Elizabeth C O’Quinn, Elizabeth Hill, Thao Vu, Debashis Ghosh, Anand S Mehta, Kristin Wallace, and Alexander V Alekseyenko. Spaceanova: Spatial co-occurrence analysis of cell types in multiplex imaging data using point process and functional anova. Journal of Proteome Research, 23(4):1131–1143, 2024.

[24] Monica T Dayao, Alexandro Trevino, Honesty Kim, Matthew Ruffalo, H Blaize D’Angio, Ryan Preska, Umamaheswar Duvvuri, Aaron T Mayer, and Ziv Bar-Joseph. Deriving spatial features from in situ proteomics imaging to enhance cancer survival analysis. Bioinformatics, 39(Supplement 1):i140–i148, 2023.

[25] Herbert Edelsbrunner and John L Harer. Computational topology: an introduction. American Mathematical Society, 2022.

[26] Oliver Vipond, Joshua A Bull, Philip S Macklin, Ulrike Tillmann, Christopher W Pugh, Helen M Byrne, and Heather A Harrington. Multiparameter persistent homology landscapes identify immune cell spatial patterns in tumors. Proceedings of the National Academy of Sciences, 118(41):e2102166118, 2021.

[27] Katherine Benjamin, Aneesha Bhandari, Jessica D Kepple, Rui Qi, Zhouchun Shang, Yanan Xing, Yanru An, Nannan Zhang, Yong Hou, Tanya L Crockford, et al. Multiscale topology classifies cells in subcellular spatial transcriptomics. Nature, pages 1–7, 2024.

[28] Andrew Aukerman, Mathieu Carrière, Chao Chen, Kevin Gardner, Raúl Rabadán, and Rami Vanguri. Persistent homology based characterization of the breast cancer immune microenvironment: A feasibility study. Journal of Computational Geometry, 12(2):183–206, 2022.

[29] Lorin Crawford, Anthea Monod, Andrew X Chen, Sayan Mukherjee, and Raúl Rabadán. Predicting clinical outcomes in glioblastoma: an application of topological and functional data analysis. Journal of the American Statistical Association, 115(531):1139–1150, 2020.

[30] Michael C Wu, Seunggeun Lee, Tianxi Cai, Yun Li, Michael Boehnke, and Xihong Lin. Rare-variant association testing for sequencing data with the sequence kernel association test. The American Journal of Human Genetics, 89(1):82–93, 2011.

[31] Ni Zhao, Jun Chen, Ian M Carroll, Tamar Ringel-Kulka, Michael P Epstein, Hua Zhou, Jin J Zhou, Yehuda Ringel, Hongzhe Li, and Michael C Wu. Testing in microbiome-profiling studies with mirkat, the microbiome regression-based kernel association test. The American Journal of Human Genetics, 96(5):797–807, 2015.

[32] Tusharkanti Ghosh, Victor Lui, Pratyaydipta Rudra, Souvik Seal, Thao Vu, Elena Hsieh, and Debashis Ghosh. The cytokernel user’s guide. dim, 126:8, 2022.

[33] Nina Otter, Mason A Porter, Ulrike Tillmann, Peter Grindrod, and Heather A Harrington. A roadmap for the computation of persistent homology. EPJ Data Science, 6:1–38, 2017.

[34] Dawei Liu, Xihong Lin, and Debashis Ghosh. Semiparametric regression of multidimensional genetic pathway data: Least-squares kernel machines and linear mixed models. Biometrics, 63(4):1079–1088, 2007.

[35] Yaowu Liu and Jun Xie. Cauchy combination test: a powerful test with analytic p-value calculation under arbitrary dependency structures. Journal of the American Statistical Association, 2019.

[36] Alex Soupir, Christopher Wilson, Jordan Creed, Julia Wrobel, Oscar Ospina, and Brooke Fridley. scSpatialSIM: A Point Pattern Simulator for Spatial Cellular Data, 2023. R package version 0.1.3.3.

[37] Ton N Schumacher and Daniela S Thommen. Tertiary lymphoid structures in cancer. Science, 375(6576):eabf9419, 2022.

[38] Eric Berry, Yen-Chi Chen, Jessi Cisewski-Kehe, and Brittany Terese Fasy. Functional summaries of persistence diagrams. Journal of Applied and Computational Topology, 4(2):211–262, 2020.

[39] Raoul R Wadhwa, Drew FK Williamson, Andrew Dhawan, and Jacob G Scott. Tdastats: R pipeline for computing persistent homology in topological data analysis. Journal of open source software, 3(28):860, 2018.

[40] Tianxi Cai, Giulia Tonini, and Xihong Lin. Kernel machine approach to testing the significance of multiple genetic markers for risk prediction. Biometrics, 67(3):975–986, 2011.

[41] Anna Plantinga, Xiang Zhan, Ni Zhao, Jun Chen, Robert R Jenq, and Michael C Wu. Mirkat-s: a community-level test of association between the microbiota and survival times. Microbiome, 5:1–13, 2017.

[42] Bradley Efron. The efficiency of cox’s likelihood function for censored data. Journal of the American statistical Association, 72(359):557–565, 1977.

43. Robert B Davies. The distribution of a linear combination of *χ*2 random variables. Journal of the Royal Statistical Society Series C: Applied Statistics, 29(3):323–333, 1980.

