## Supplementary Materials for "Detecting Clinically Relevant Topological Structures in Multiplexed Spatial Proteomics Imaging Using TopKAT"

#### 1 Small Sample Correction for Survival Outcomes

The test described in Section 6.4 of the main manuscript is conservative for small sample sizes. As such, we use a correction for small sample sizes as described in Plantinga et al. (2017) [1]. We consider modifying the variance-component score test statistic:

$$Q^k = \frac{\hat{\mathbf{m}}^T \mathbf{K}^k \hat{\mathbf{m}}}{\hat{\mathbf{m}}^T \hat{\mathbf{m}}} \tag{1}$$

This modified test statistic has been shown to reduce the conservativeness of the kernel association test for small sample sizes and complicated kernels in microbiome analyses [1]. More details and derivations of this test statistic can be found in Plantinga et al. (2017). In Supplementary Materials Sections 3 and 4 we describe analogous small sample corrections for continuous and binary outcomes, respectively.

### 2 Multiple Kernels

In the two-dimensional imaging context, we have two kernel matrices, $\mathbf{K}^0$ and $\mathbf{K}^1$, corresponding to degree-0 and degree-1 homologies. We propose to construct an aggregate kernel matrix comprised of a weighted linear combination of $\mathbf{K}^0$ and $\mathbf{K}^1$. We vary the weights, $\omega$, across a range of values between 0 and 1 and combine the p-values resulting from testing $H_0$ using the Cauchy combination test [2]. The weighted combination of kernel matrices is:

$$\mathbf{K} = (1-\omega)\mathbf{K}^0 + \omega\mathbf{K}^1 \tag{2}$$

The resulting p-values are denoted $p_1, \ldots, p_\Omega$ where $\omega = 1, \ldots, \Omega$. Using the Cauchy combination test, which is appropriate for combining p-values with an arbitrary dependence structure, we can obtain an omnibus p-value, $p^{omnibus}$. To apply the Cauchy combination test, we construct a weighted aggregation of the transformed p-values which yields the following test statistic:

$$T = \sum_{\omega=1}^{\Omega} c_\omega \tan\left[\pi(0.5 - p_\omega)\right] \tag{3}$$

where $\sum_{\omega=1}^{\Omega} c_\omega = 1$. For simplicity, we fix $c_\omega = 1/\Omega$ for all $\omega$. Then, $T \sim \text{Cauchy}(0,1)$ from which we obtain $p^{omnibus}$. This p-value describes the significance of the association between persistence diagrams and the log-hazard for the event in the context of survival, the average response for a continuous outcome, and the log-odds of the event given a binary outcome.

### 3 TopKAT for Continuous Outcomes

We now describe the test statistic and small sample correction used when testing for an association between persistence diagrams and continuous patient-level outcomes, following the discussion provided in Zhao et al. (2015) [3]. For a continuous outcome, our kernel machine regression model can be represented as:

$$\mathbf{y} = \beta_0 \mathbf{1} + \boldsymbol{\beta}\mathbf{X} + f(\mathbf{Z}) + \boldsymbol{\epsilon} \tag{4}$$

where $\mathbf{1} : n \times 1$ is a vector of 1s, $\mathbf{X} : n \times p$ is a matrix of covariates to adjust for, $f(\cdot)$ is an unknown function, and $\mathbf{Z}$ is a vector of $n$ persistence diagrams. As described in the main manuscript, $f(\cdot)$ is assumed to fall within a space spanned by a positive definite kernel $K(\cdot, \cdot)$. We can view $f(\mathbf{Z})$ is a sample-level random effect through the relationship between kernel machine regression and linear mixed modeling [4]. $f(\mathbf{Z})$ is assumed to have mean 0 and covariance matrix equal to $\tau\mathbf{K}$. We can then test for an effect of $f(\mathbf{Z})$ on $\mathbf{y}$ by equivalently testing if $\tau = 0$. In other words, our null and alternative hypotheses are $H_0 : \tau = 0$ vs. $H_1 : \tau > 0$. The corresponding variance component score test statistic for $H_0$ is:

$$Q = \frac{1}{2\hat{\sigma}_0^2}(\mathbf{y} - \hat{\mathbf{y}}_0)^T \mathbf{K}(\mathbf{y} - \hat{\mathbf{y}}_0) \tag{5}$$

where $\hat{\mathbf{y}}_0$ is the fitted value for $\mathbf{y}$ under $H_0$ and $\hat{\sigma}_0^2$ is the estimated error variance under $H_0$.

Testing using $Q$ can be conservative with small sample sizes due to uncertainty in estimation of $\sigma_0^2$. Accounting for this uncertainty may increase power. To this end, we use the proposed test statistic given in Chen et al. (2016) [5] which is a modification of $Q$ given in Equation 5. Let $\mathbf{P}_0 = \mathbf{I} - \mathbf{X}(\mathbf{X}^T\mathbf{X})^{-1}\mathbf{X}^T$. The small-sample corrected test statistic is:

$$Q = \frac{\boldsymbol{\epsilon}^T \mathbf{P}_0 \mathbf{K} \mathbf{P}_0 \boldsymbol{\epsilon}}{\boldsymbol{\epsilon}^T \mathbf{P}_0 \boldsymbol{\epsilon}} \tag{6}$$

where $\boldsymbol{\epsilon} = \mathbf{y} - \hat{\mathbf{y}}$.

### 4 TopKAT for Binary Outcomes

The kernel machine regression model for a binary outcome can be written as:

$$\text{logit}\,[P(\mathbf{y}=1)] = \beta_0\mathbf{1} + \mathbf{X}\boldsymbol{\beta} + f(\mathbf{Z}) \tag{7}$$

We can again test for the effect of $f(\mathbf{Z})$ on $\text{logit}\,[P(\mathbf{y}=1)]$ by leveraging the relationship between kernel machine regression and generalized linear mixed modeling. This equates testing for the effect of $f(Z)$ as testing $H_0 : \tau = 0$ and $H_1 : \tau > 0$ where $\tau$ is as defined in Section 3. We can derive the following variance component score test statistic as:

$$Q = \frac{1}{2}(\mathbf{y} - \hat{\mathbf{y}}_0)^T \mathbf{K} (\mathbf{y} - \hat{\mathbf{y}}_0) \tag{8}$$

where $\hat{\mathbf{y}}_0$ is the fitted probability of $P(\mathbf{y}=1)$ under $H_0 : \tau = 0$.

To address the conservativeness of this test as a result of small sample sizes or deviations from the assumed model structure (i.e. due to overdispersion), we can use the modified test statistic proposed by Chen et al. (2016) [5] for binary outcomes in kernel machine regression. The modified test statistic is based on the iteratively reweighted least squares (IRLS) algorithm for estimating the parameters in a logistic regression model accounting for overdispersion. Define $\mu_i = P(y_i = 1)$ and $\text{logit}(\boldsymbol{\mu}) = \mathbf{X}\boldsymbol{\gamma}$. The modified test statistic is:

$$Q = \frac{(\tilde{\mathbf{y}} - \mathbf{X}\tilde{\gamma})^T \mathbf{DKD} (\tilde{\mathbf{y}} - \mathbf{X}\tilde{\gamma})}{\hat{\phi}} \tag{9}$$

where $\tilde{\mathbf{y}}$ is the working response at convergence of the IRLS algorithm, $\mathbf{D}$ is a diagonal matrix with diagonal elements equal to $\tilde{\mu}_i(1 - \tilde{\mu}_i)$, and $\hat{\phi}$ is the IRLS estimate of the overdispersion parameter.

### 5 Simulation Study

#### 5.1 Simulation Set-up

To evaluate the performance of TopKAT, we considered a simulation study in which we simulated images exhibiting different point patterns and geometric structures. As described in the main manuscript, we simulated $n = 100$ samples each of dimension $100 \times 100$. These images were split into two groups of size 50 and for each group we chose a different geometric structure to generate. We considered simulating a random number (between two and six) of squares ("square"), a random number (between two and six) of loops ("loop"), two bivariate Gaussian distributions ("clusters"), simulated real tissue using the `scSpatialSIM R` package ("real tissue") [6], and complete spatial randomness ("CSR"). In simulating "real" tissue, we used the `scSpatialSIM` package to generate two kinds of tissue with intensity $\lambda = 0.2$. The package simulates cell locations within a Gaussian kernel. We fixed the minimum standard deviation of the kernel at 0.5 and the maximum at 25. We then split the two tissue types to represent our two groups. We denote each condition in our simulation based on the geometric shape chosen to be simulated for each group: (1) square vs. CSR, (2) loop vs. CSR, (3) square vs. loop, (4) cluster vs. CSR, and (5) simulated real tissue. In Section 5.2 of the Supplementary Materials, we exhibit example images of each geometric shape. We then compared TopKAT to SPOT [7], SPF [8], and FunSpace [9] across the five conditions.

We considered simulating a survival outcome. Since the samples were split into two groups, we simulated the outcomes from two different distributions. We generated the survival times from an exponential distribution with rates either equal to 0.2 or 0.1 for a hazard ratio of 2. We randomly censored 10% of events in each group.

We considered TopKAT across a series of weights, $\omega$, that we aggregated kernel matrices across. We present results when considering only connected components, only loops, and aggregating across three weighted aggregations of the kernel matrices for $\omega = 0, 0.5, 1$. We compare TopKAT to SPOT [7], SPF [8], and FunSpace [9]. These methods all describe the spatial distribution of points in the image using the spatial point process model. SPOT computes a spatial summary statistic for a range of radii (here fixed between $0, 1, \ldots, 250$). The association between the spatial summary statistic and outcome is tested at each radius and the resulting p-values are aggregated using the Cauchy combination test [2]. In the case of a survival outcome, the p-values arise from a Wald test for $\beta_r = 0$ where $\beta_r$ corresponds to the regression coefficient for the effect of the spatial summary statistic evaluated at radius $r$ on the log-hazard for the event. SPF and FunSpace are both functional analytic approaches to describing the spatial distribution of cells. Both evaluate a spatial summary statistic across a range of radii (here, also between $0, 1, \ldots, 250$). The resulting curve is treated as a functional covariate in a regression model in the case of SPF. In the case of FunSpace, the curves across samples are decomposed using functional principal components analysis and the scores are treated as fixed covariates in the appropriate regression model for the outcome.

#### 5.2 Example Simulation Images

Figure 1 illustrates example images simulated throughout the simulation study.

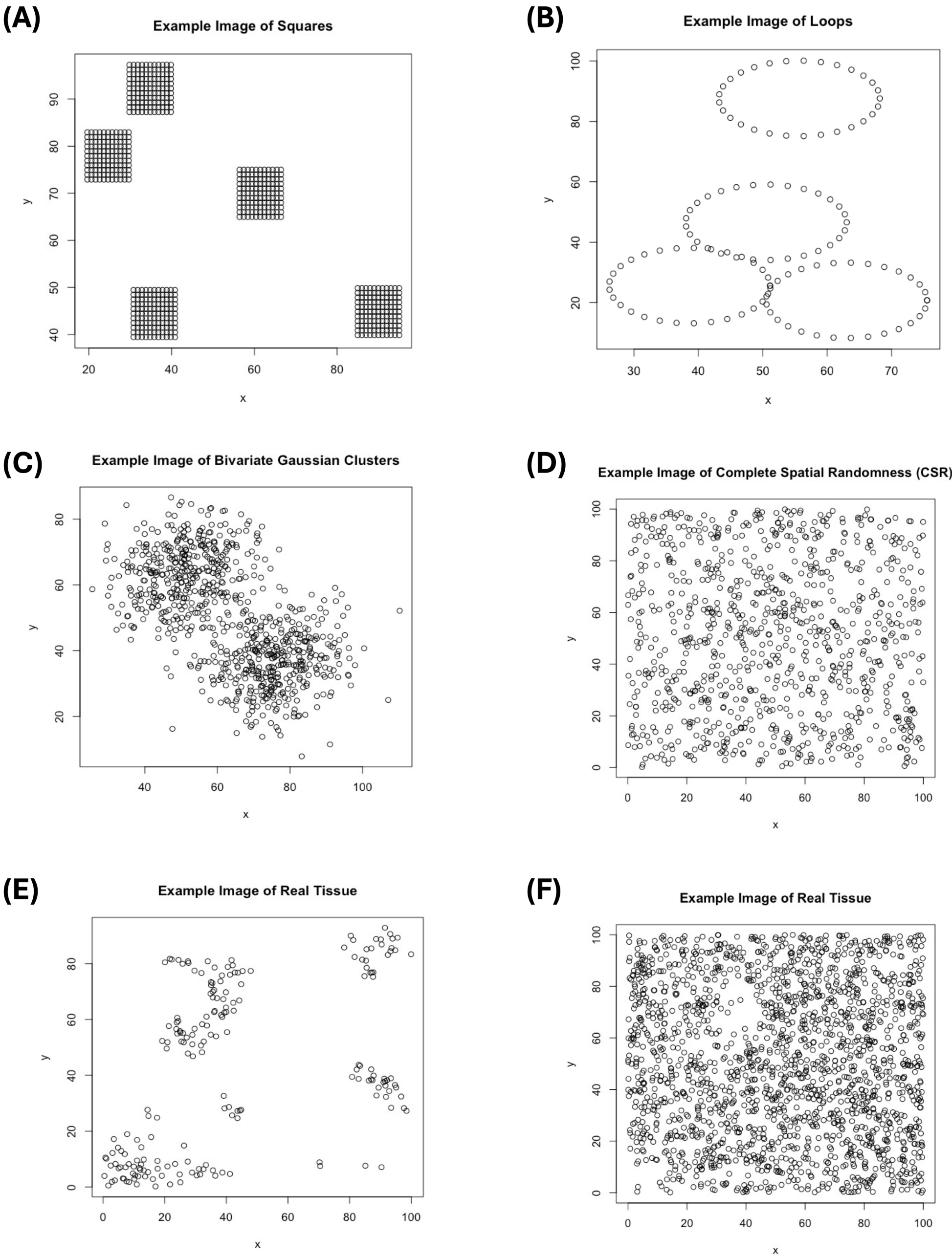


Figure 1: Examples of spatial arrangements of cells simulated during our simulation study

### 6 More Details on Multiplexed Ion Beam Imaging in Triple Negative Breast Cancer

Here we describe additional details on the application of TopKAT to multiplexed ion beam imaging time-of-flight (MIBI-TOF) data from a study of triple negative breast cancer [10]. Each MIBI-TOF image was of dimension $2048 \times 2048$ so we constructed a Rips filtration for each image with a maximum diameter of 2048. We computed the distance between the persistence diagrams using the distance measure provided in Equation 3 in the main manuscript. We then converted these pairwise distance matrices into kernel matrices using the formula in Equation 4 in the main manuscript. We aggregated the p-values across a weighted combination of kernel matrices using weights $\omega = (0, 0.5, 1)$.

We first examined whether persistent homology could characterize the topological differences between TME categories, which were provided in the published data. We conducted pairwise comparisons between mixed vs. compartmentalized, mixed vs. cold, and compartmentalized vs. cold samples to examine if TopKAT could distinguish between their topological profiles. For each pair, we treated the TME categories as a binary outcome and computed a p-value using the kernel machine regression framework described in Section 4 of the Supplementary Materials. The TopKAT p-values for each pairwise comparison were as follows: mixed vs. compartmentalized ($p = 0.0005$), mixed vs. cold ($p = 0.0126$), compartmentalized vs. cold ($p = 0.00005$). This suggests persistent homology can capture the topological differences between these TMEs. Mixed vs. cold samples are more challenging to distinguish, as evidenced by a higher (though still significant) TopKAT p-value. This is because the main distinction between these samples is the distance between cells and not the presence or absence of large loops. Mixed vs. compartmentalized samples, however, differ in both distance between cells and the presence of large holes (loops) among the immune cells.

In Section 2.3 of the main manuscript, we described identifying the distance at which mixed vs. compartmentalized samples showed the most significant difference in topological structure. In Figure 2, we show the relationship between distance used in the Rips filtration and TopKAT p-value. The most significant distance, $2\epsilon = 73.6$, is identified by a vertical dashed line.

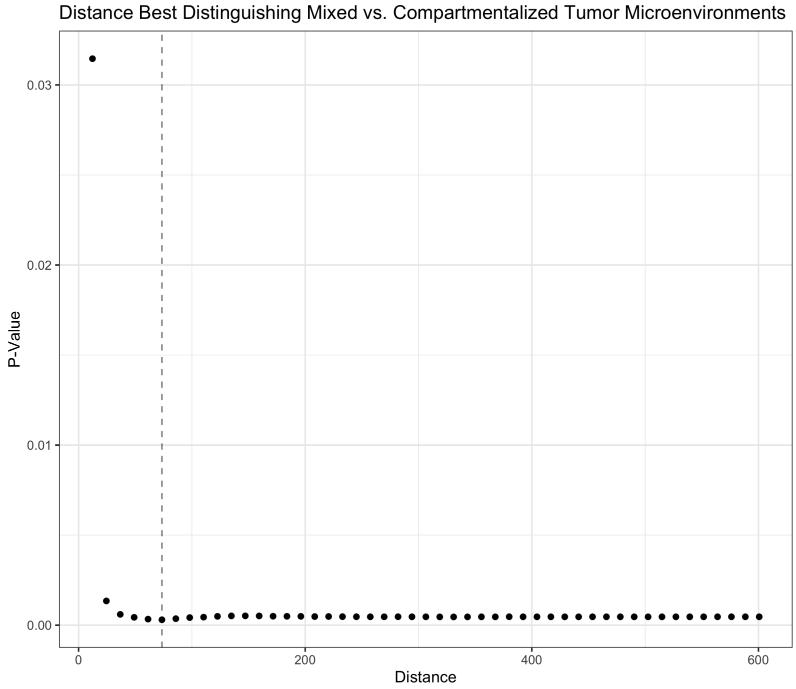


Figure 2: The significance of each distance in describing the association between the topological structure of each image and compartmentalized vs. mixed tumor microenvironments (TMEs) in a study of triple negative breast cancer. The most significant distance, the distance that best distinguished these two classes of TMEs, is identified with a vertical dashed line.

### 7 Application to Immune Checkpoint Blockade Study in Triple Negative Breast Cancer

We applied TopKAT to imaging mass cytometry data from the NeoTRIP study which was also a study in triple negative breast cancer [11]. NeoTRIP compared neoadjuvant chemotherapy plus immunotherapy (immune checkpoint blockade, or ICB) to neoadjuvant chemotherapy. Investigators were interested in how the TME was altered following ICB in patients who responded to treatment compared to those who did not.

In this analysis, we focused on the TMEs of patients in the chemotherapy plus ICB treatment arm. We examined the geometric structures among tumor, immune, and endothelial cells in the tumor biopsies after treatment to examine how the TME differed between responders, who exhibited pathological complete response (pCR), and non-responders, who experienced recurrent disease (RD). We treated pCR vs. RD as a binary outcome to examine the differences among topological structures between these groups. Some patients within this treatment arm and at the post-treatment timepoint had more than one image available, so we selected one image from each person that contained the most cells. We used the kernel machine regression framework described in Section 4 of the Supplementary Materials to compare pCR and RD samples. This dataset contained 106 tumor biopsies post-treatment on the chemotherapy plus ICB treatment arm. Of these, 57 patients experienced pCR and 49 experienced RD. We applied our analysis to all detected cell types found across TMEs and used weights $\omega = (0, 0.5, 1)$ to aggregate p-values across different weighting schemes of the connected component- and loop-kernel matrices.

The resulting p-value for the association with likelihood of pathologic complete response was $p = 0.0075$, suggesting patients who experienced the same treatment outcomes also had TMEs which showcased similar topological structure among the immune cells after treatment. Example persistence diagrams for a sample which exhibited pCR and a sample which exhibited RD are shown in Figures 4A and 4B. pCR TMEs had, on average, more connected components (1956.1) vs. RD (1093.1) and more loops (676.0) vs. RD (296.1). The average maximum lifespan, however, for connected components was shorter among pCR TMEs (94.4 vs. 103.4) but longer for loops among

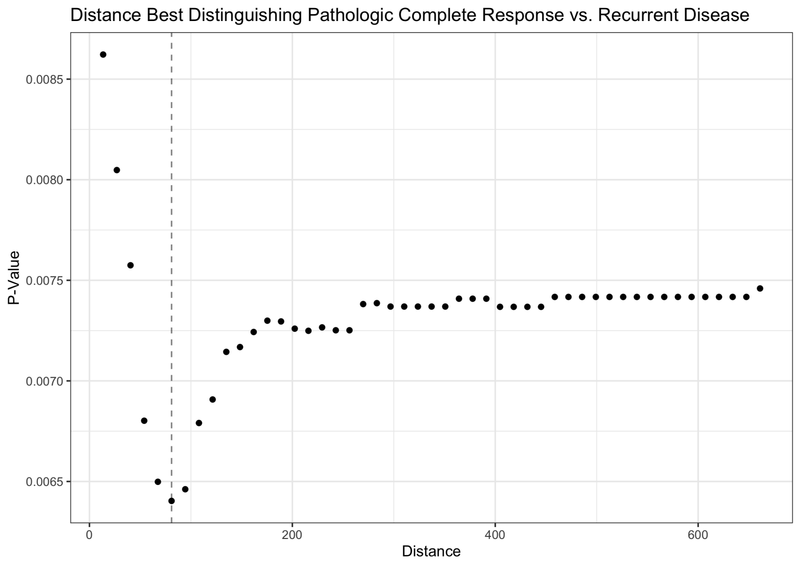


Figure 3: Illustrating the distance between cells at which samples corresponding to patients exhibiting pathologic complete response vs. those corresponding to patients exhibiting recurrent disease are most distinct topologically. This occurs at distance $2\epsilon = 80.9$. We then illustrate the average number of connections between cell types at this distance in Figures 5A and 5B.

pCR TMEs (86.1 vs. 78.8).

We then examined the distance at which pCR and RD TMEs showed the most significant topological differences (Figure 3). We found that a distance of $2\epsilon = 80.9$ produced the most significant difference between TMEs corresponding to patients exhibiting pCR vs. RD (Figures 4C and 4D). At this distance, we observed acute differences in the connections among immune cells. In RD TMEs, CD8 T cells, CD20+ B cells, and CD4 T cells were all highly connected, with few connections among other cell types. In pCR TMEs, in contrast, these cell types were highly connected, as were fibroblasts, mcyofibroblasts, M2 macrophages, and dendritic cells (DCs).

We compared the results using TopKAT to SPOT, SPF, or FunSpace which were considered in the Simulation Study described in Section 2.2 of the main manuscript. Given a range of radii between 0 and 72, we did not find significant associations between the spatial arrangement of immune cells and the log-odds of pCR using these methods.

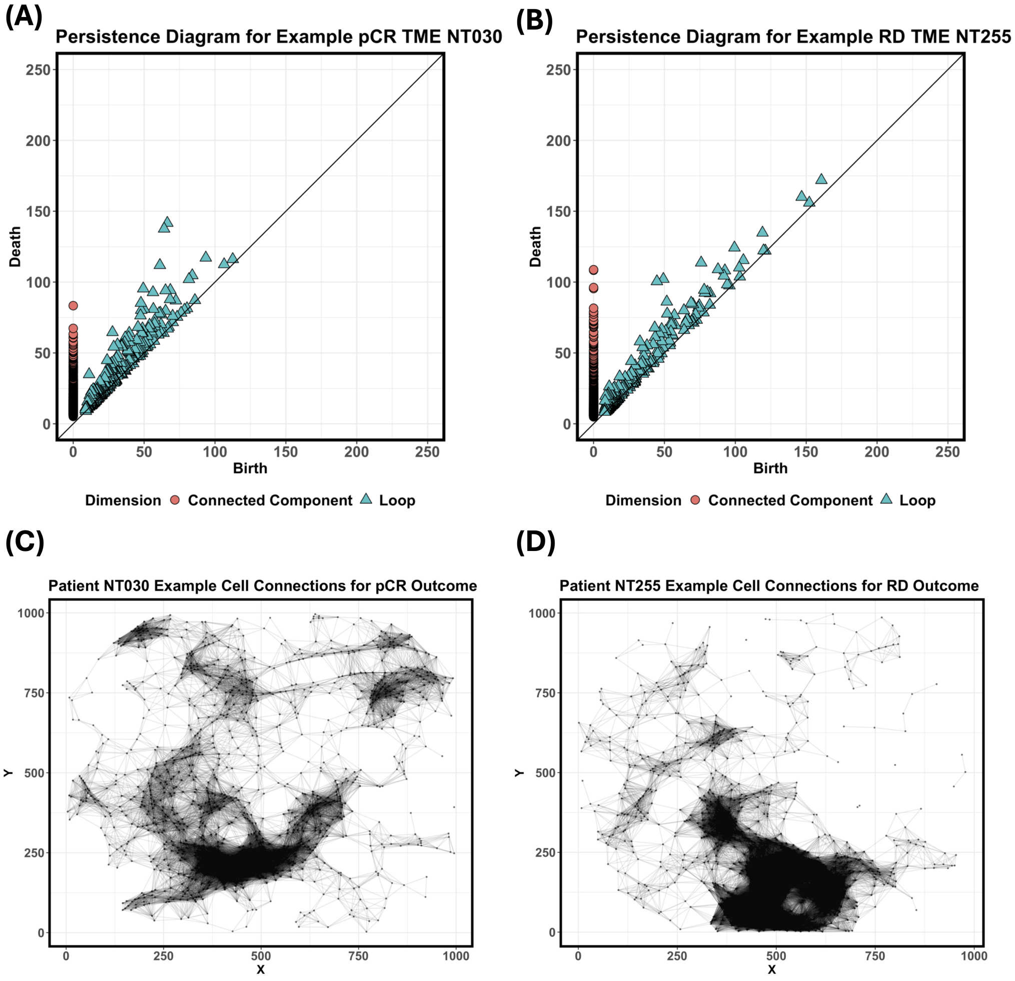


Figure 4: We illustrate the topological structure in two example samples corresponding to patients exhibiting either pathologic complete response (pCR) or recurrent disease (RD). Figure 4A shows the persistence diagram corresponding to an example sample from a patient exhibiting pCR. Figure 4B shows the persistence diagram corresponding to an example sample corresponding to a patient exhibiting RC. pCR samples exhibited fewer connected components and loops among cells. However, the lifespans for these features in pCR samples were shorter, on average, than in RD samples. Figure 4C shows the connectivity among the cells at the best distance given in Figure 3 in the example pCR sample. Figure 4D shows the connectivity among cells for the same distance in the example RD sample.

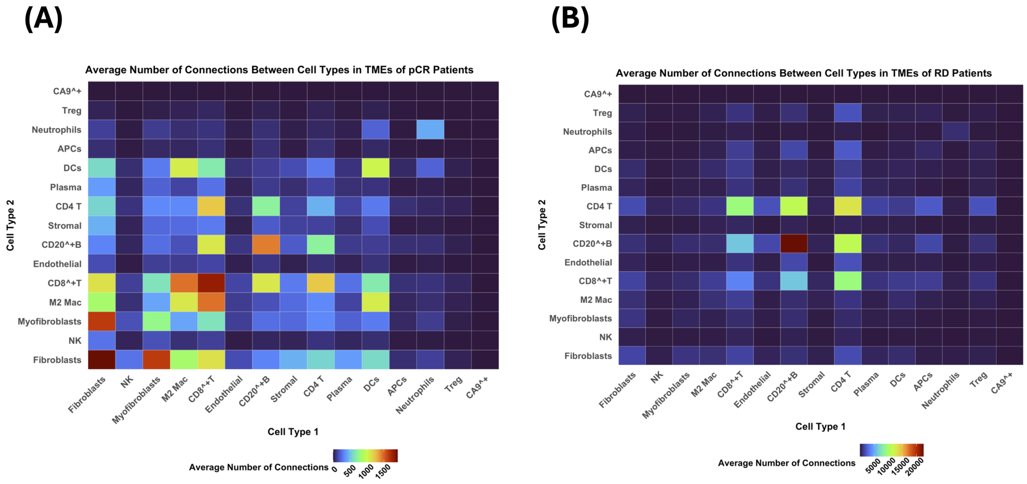


Figure 5: Average connectivity among cells at a clinically-relevant distance in samples treated with chemotherapy plus immunotherapy. Figure 5A shows the average connectivity among immune cells in patients that showed pathologic complete response (pCR) or patients who responded to treatment. Figure 5B shows the average connectivity among immune cell in patients that showed recurrent disease (RD) or patients who did not respond to treatment.

### References


[1] Anna Plantinga, Xiang Zhan, Ni Zhao, Jun Chen, Robert R Jenq, and Michael C Wu. Mirkat-s: a community-level test of association between the microbiota and survival times. *Microbiome*, 5:1–13, 2017.

[2] Yaowu Liu and Jun Xie. Cauchy combination test: a powerful test with analytic p-value calculation under arbitrary dependency structures. *Journal of the American Statistical Association*, 2019.

[3] Ni Zhao, Jun Chen, Ian M Carroll, Tamar Ringel-Kulka, Michael P Epstein, Hua Zhou, Jin J Zhou, Yehuda Ringel, Hongzhe Li, and Michael C Wu. Testing in microbiome-profiling studies with mirkat, the microbiome regression-based kernel association test. *The American Journal of Human Genetics*, 96(5):797–807, 2015.

[4] Dawei Liu, Xihong Lin, and Debashis Ghosh. Semiparametric regression of multidimensional

genetic pathway data: Least-squares kernel machines and linear mixed models. *Biometrics*, 63(4):1079–1088, 2007.

[5] Jun Chen, Wenan Chen, Ni Zhao, Michael C Wu, and Daniel J Schaid. Small sample kernel association tests for human genetic and microbiome association studies. *Genetic epidemiology*, 40(1):5–19, 2016.

[6] Alex Soupir, Christopher Wilson, Jordan Creed, Julia Wrobel, Oscar Ospina, and Brooke Fridley. *scSpatialSIM: A Point Pattern Simulator for Spatial Cellular Data*, 2023. R package version 0.1.3.3.

[7] Sarah N Samorodnitsky, Katie M Campbell, Antoni Ribas, and Michael C Wu. A spatial omnibus test (spot) for spatial proteomic data. *bioRxiv*, pages 2024–03, 2024.

[8] Thao Vu, Julia Wrobel, Benjamin G Bitler, Erin L Schenk, Kimberly R Jordan, and Debashis Ghosh. Spf: a spatial and functional data analytic approach to cell imaging data. *PLoS computational biology*, 18(6):e1009486, 2022.

[9] Thao Vu, Souvik Seal, Tusharkanti Ghosh, Mansooreh Ahmadian, Julia Wrobel, and Debashis Ghosh. Funspace: A functional and spatial analytic approach to cell imaging data using entropy measures. *PLOS Computational Biology*, 19(9):e1011490, 2023.

[10] Leeat Keren, Marc Bosse, Diana Marquez, Roshan Angoshtari, Samir Jain, Sushama Varma, Soo-Ryum Yang, Allison Kurian, David Van Valen, Robert West, et al. A structured tumor-immune microenvironment in triple negative breast cancer revealed by multiplexed ion beam imaging. *Cell*, 174(6):1373–1387, 2018.

[11] Xiao Qian Wang, Esther Danenberg, Chiun-Sheng Huang, Daniel Egle, Maurizio Callari, Begoña Bermejo, Matteo Dugo, Claudio Zamagni, Marc Thill, Anton Anton, et al. Spatial predictors of immunotherapy response in triple-negative breast cancer. *Nature*, 621(7980):868–876, 2023.
